# Glyoxal Does Not Preserve Cellular Proteins as Accurately as PFA: A Microscopical Survey of Epitopes

**DOI:** 10.1101/625863

**Authors:** Ferda Topal Celikkan, Ceren Mungan, Merve Sucu, Fatma Uysal, Selda Kahveci, Serhat Hayme, Nilay Kuscu, Sinan Ozkavukcu, Ciler Celik-Ozenci, Alp Can

## Abstract

Chemical fixation is one of the most critical steps to retaining cellular targets as naturally as possible. Recent developments in microscopy allow sophisticated detection and measuring techniques with which spatio-temporal molecular alterations is conceivable. Here, we document the fixation competence of glyoxal (Gly), a less-toxic dialdehyde molecule, and paraformaldehyde (PFA) side-by-side (with or without Triton-X 100 permealization) in live- and fixed-cell preparations including human stem cells, spermatozoa, mouse oocytes/embryos using super-resolution microscopy. Although Gly seemed to act faster than PFA, catastrophic consequences were found not acceptable, especially in oocytes and embryos. Due to cell lysate and immunocytochemistry surveys, it was obvious that PFA is superior to Gly in retaining cellular proteins *in situ* with little/no background staining. In many samples, PFA revealed more reliable and consistent results regarding the protein quantity and cellular localization corresponding to previously defined patterns in the literature. Although the use of Gly is beneficial as indicated by previous reports, we concluded that it does not meet the requirement for proper fixation, at least for the tested cell types and proteins.

## Introduction

Fixation is a fundamental and initial step in histochemical and cytochemical investigations, mainly aiming at preserving tissues, cells, and cellular components from autolytic deterioration. Microscopic structures can subsequently be observed *in situ* after suitable cell/tissue preparations and labelling procedures. For this purpose, a number of chemical fixatives have been used since the nineteenth century (Howat & Wilson, 2014). First introduced in 1893, formaldehyde (FA), as a member of aldehyde fixatives, in the form of a water-based, diluted solution of 1:10 has been the most commonly-used chemical fixative, and it is found in many fixative cocktails (Marcon, Bressenot et al., 2009). Paraformaldehyde (PFA), the polymerized form of FA, is generally favoured over FA because PFA crosslinks amino groups without altering the tertiary structure of proteins, thereby cellular epitopes remain relatively well-preserved in a successful labelling protocol with specific antibodies (Fujiwara, 1980, Leyton-Puig, Kedziora et al., 2016, Robinson & Snyder, 2004). PFA fixation has been widely adopted to preserve cell morphology for immunolabeling where the final concentration of PFA in the fixative solution is around 3-4% (Kim, Kim et al., 2017). Nonetheless, PFA has also been associated with various problems, ranging from loss of antigenicity to changes in morphology during fixation.

Glyoxal (Gly), the smallest dialdehyde reagent with a structural formula that resembles two formaldehyde molecules bonded back-to-back, has also been tested as a fixative since the early 1960s (Sabatini, Bensch et al., 1963), albeit in fewer studies. Due to a better safety profile, faster reaction rate, and selective control over crosslinking, Gly is considered to retain immunoreactivity and decreases the need for antigen retrieval (Dapson, 2007). With suitable catalysts or other reaction accelerators, Gly forms two-carbon adducts with nearly all end groups in proteins and carbohydrates, leaving most of them unimpaired for subsequent immunohistochemical (IHC) demonstration. Due to the previously reported disadvantages of PFA, Gly has recently been suggested superior to PFA as being less toxic, and preserves cells better as investigated using super-resolution microscopy (Richter, Revelo et al., 2018). There are also a number of reports comparing the staining and IHC labelling results between Gly- and FA-fixed tissues. PreFer, a commercial form of Gly, was found to be unsuitable because of poor preservation of tissue morphology (Atkins, Reiffen et al., 2004); Compared with FA, Gly was found better at preserving the cell membranes and nuclear chromatin in paraffin-embedded tissue sections (Dapson R. W., 2006); Oestrogen receptor staining with antigen retrieval was found significantly weaker in the Gly-fixed specimens than in FA-fixed specimens (Umlas & Tulecke, 2004). No significant differences were reported between FA and Gly fixation in histomorphometry (Wang, Lee et al., 2011); IHC and Western blot (WB) (Paavilainen, Edvinsson et al., 2010), and fluorescence in situ hybridization (Bussolati, Annaratone et al., 2017) assays.

Besides fixation, many samples require a detergent extraction step for exposing antigenic sites to antibodies. As such, using a non-ionic detergent such as Triton X-100 (TX) may be an essential step for improving the penetration of the antibody. TX effectively solubilizes complexes such as biologic membranes, and it apparently does not inhibit the antigen-antibody reaction (Dimitriadis, 1979). However, it is not appropriate for membrane-associated antigens because it destroys lipids among membranous proteins. TX may be added to fixation solutions, thus simultaneously performing fixation and extraction actions. Alternatively, in many settings, TX extraction is applied following cell fixation in daily use.

In this study, we aimed to assess whether Gly could provide a successful and consistent cell fixation in favour of PFA in various cell preparations and types such as cultured human stromal/stem cells, human spermatozoa smears, isolated mouse granulosa-enclosed or naked oocytes and embryos, all of which were analysed during (live-cell imaging) and after fixation using a wide-range of antibodies raised against diverse cellular proteins. The study groups were designed to test the PFA (3.5%) and Gly (3%) alone, as well as with the addition of TX (0.1%). Additionally, TX (0.1%) application was also performed after the samples were fixed.

In short, PFA was found to be more consistent and potent as a fixative compared with Gly in many circumstances. Here, we present the relatively poor fixation capacity of Gly, proven by a series of quantitative and qualitative data, some of which were obtained using super-resolution microscopy. The overall results seemed quite different from previously published data. Interestingly, however, PFA alone with no addition of TX displayed a significant cytoplasmic loss by generating membrane blebs during fixation. In this article, we also demonstrated novel *in situ* remodelling of some proteins, such as the ACRBP in human spermatozoa, which have not been presented in previous literature.

## Results

### 1) Live-cell Imaging of mEmbryos and hUC-MSCs

Freshly-isolated mouse embryos (mEmbryos) and cultured human umbilical cord-derived mesenchymal stem cell (hUC-MSC) monolayers in glass-bottomed Petri dishes were microscopically examined using a laser-illuminated differential interference contrast (DIC) imaging to observe the cell dynamics during fixation. Prior to fixation, live samples in culture media were recorded in a time-lapse manner for 15-25 min (**Fig 1** and **Extended View Content**). Subsequently, the same samples were directly taken to fixation simply by replacement of the culture media with the fixative solution. As seen in **Fig 1, Extended View Content** and **Table 3**, a series of significant changes were noted during the fixation interval (20 min for mEmbryos or 15 min for hUC-MSCs) followed by a 5-min TX incubation.

**Fig 1.**
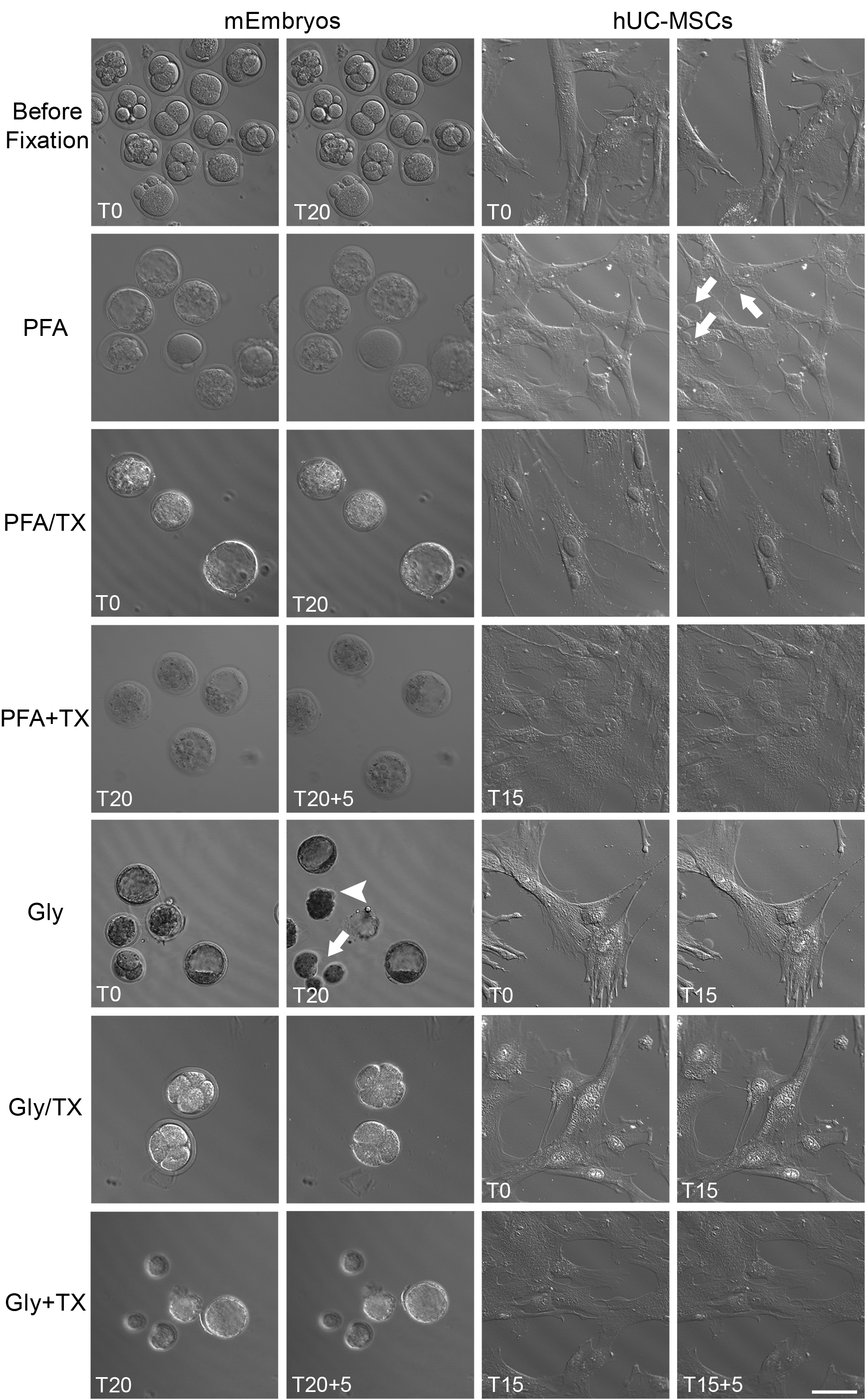
Still images before and during PFA- and Gly-containing fixative cocktails. **First row.** Live mEmbryos and hUC-MSCs before fixation. **Second-seventh rows.** mEmbryos (n=49) and hUC-MSCs (n=14) during PFA, PFA/TX, PFA+TX, Gly, Gly/TX and Gly+TX fixation procedures. Note the ZP thinning (arrowheads) and blastomere dissociation (arrow) in Gly group; cytoplasmic blebs (arrows) in PFA group; See, also **Table 3** and **Extended View Content** for live videos. Data information: T refers to time of fixation (20 min for mEmbryos; 15 min for hUC-MSC); additional 5 min TX incubation was applied in +TX groups. Scale bars: 50 μm.

**Table 1:** Fixation/Extraction protocol and experiment groups.

| Fixation/<br>Extraction<br>Agent | PFA | PFA/TX | PFA+TX | Gly | Gly/TX | Gly+TX |
| --- | --- | --- | --- | --- | --- | --- |

| **Formula** | 3.5% PFA | 3.5% PFA<br>with TX<br>(0.1%) | TX (0.1%)<br>after 3.5%<br>PFA | 3% Glyoxal | 3% Glyoxal<br>with TX<br>(0.1%) | TX (0.1%)<br>after 3%<br>Glyoxal |
| --- | --- | --- | --- | --- | --- | --- |
| **Fixation**<br>**time and**<br>**temp (°C)** | 15 min @<br>RT* | 15 min @<br>RT* | 15 min + 5<br>min @ RT* | 15 min @<br>RT* | 15 min @<br>RT* | 15 min + 5<br>min @ RT* |

\*20 minutes @ room temperature (RT) for mOocytes and mEmbryos

**Table 2.**
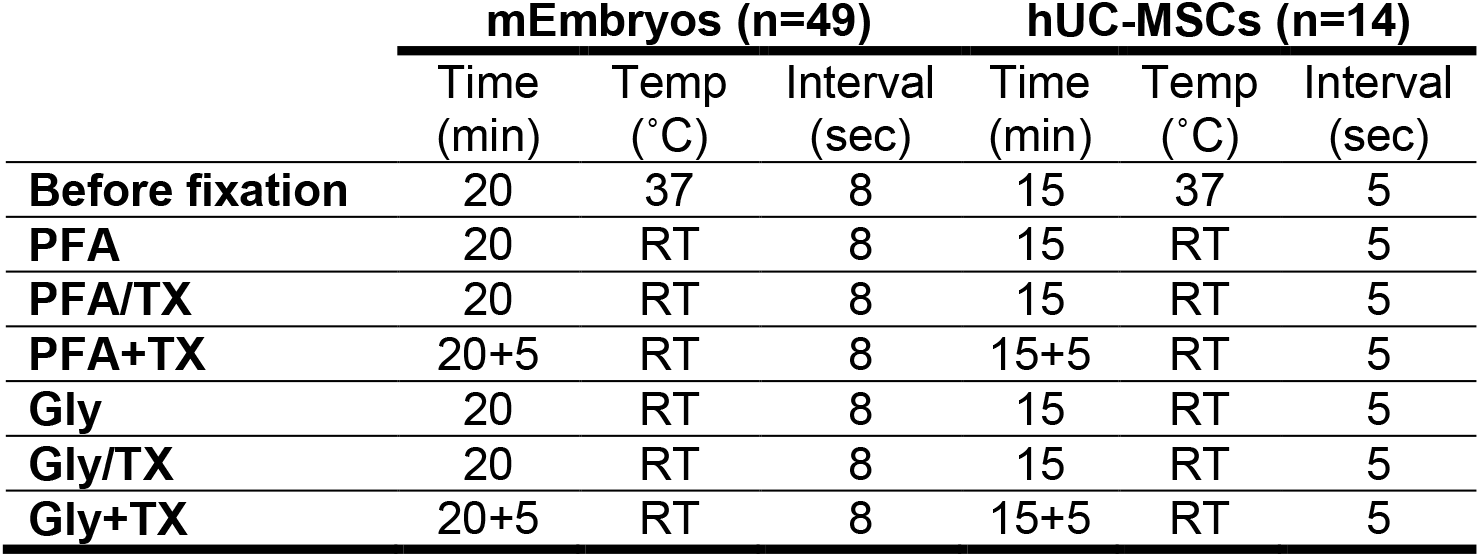
Live-cell imaging groups and protocols.

|  | **mEmbryos (n=49)** |  |  | **hUC-MSCs (n=14)** |  |  |
| --- | --- | --- | --- | --- | --- | --- |
|  | Time<br>(min) | Temp<br>(°C) | Interval<br>(sec) | Time<br>(min) | Temp<br>(°C) | Interval<br>(sec) |
| **Before fixation** | 20 | 37 | 8 | 15 | 37 | 5 |
| **PFA** | 20 | RT | 8 | 15 | RT | 5 |
| **PFA/TX** | 20 | RT | 8 | 15 | RT | 5 |
| **PFA+TX** | 20+5 | RT | 8 | 15+5 | RT | 5 |
| **Gly** | 20 | RT | 8 | 15 | RT | 5 |
| **Gly/TX** | 20 | RT | 8 | 15 | RT | 5 |
| **Gly+TX** | 20+5 | RT | 8 | 15+5 | RT | 5 |

**Table 3.** Summary of live-cell imaging results before and during fixation.

|  | **mEmbryos** | **hUC-MSCs** |
| --- | --- | --- |
| **Before fixation** | Embryos from one-cell to morula stage surrounded by an intact ZP | Normal saltatory movement of organelles and vesicles in extremely flat and widened cell bodies. Occasional retraction of cell cytoplasm (see, upper part of the Supplementary video) |
| **PFA** | Minimal thinning of ZP | Formation of large cytoplasmic blebs proceeded by plasma membrane rupture and thus loss of cytosol |
| **PFA/TX** | Swelling of blastomeres in morula and blastocysts | A minimal or no change of cell membrane integrity, extracellular flow of tiny cytoplasmic vesicles |
| **PFA+TX** | No further significant change | No further significant change |
| **Gly** | Enlargement of perivitelline space, thinning of ZP, dissolution of ZP and dissociation of blastomeres | Minimal number of tiny blebs with no sign of cytosolic loss |
| **Gly/TX** | Same effects as described in Gly fixation | No change of cell membrane integrity, no cytosolic loss |
| **Gly+TX** | No further significant change | No further significant change |

### 2) Effectiveness of PFA vs. Gly in Whole Cell Lysates

The protein cross-linking capacity of the two different fixative (PFA and Gly) were tested by evaluating the intensity of unfixed total protein bands per lane (**Fig 2A** and **Fig 3A**). To compare the efficiency of fixation, the bands that survived fixation were summed and expressed as the percentage (%) of an unfixed control. Thirty-seven per cent of proteins in human spermatozoa (hSpermatozoa) specimens remained unfixed with PFA, and 62% of proteins were found unfixed by Gly (**Fig 2A1**). Fixation of hUC-MSC lysates revealed more consistent results because PFA was not able to fix only 20% of proteins, and Gly was not able to fix 55% of proteins (**Fig 2A2**). We then tested the degree of fixation of PFA and Gly for three major cytoskeletal proteins (β-actin, vimentin and α/β-tubulin), all of which have low molecular weights (42, 57 and 50 kDa, respectively), after they were labelled with fluorescent markers. In hSpermatozoa, 0% of β-actin was unfixed with PFA, and 60% was unfixed with Gly (**Fig 2B** and **B1**); 16% of vimentin was unfixed with PFA, and 50% was unfixed with Gly (**Fig 2C** and **C1**); 8% of αβ-tubulin was unfixed with PFA, and 43% was unfixed with Gly (**Fig 2D** and **D1**). In hUC-MSCs, 17% of β-actin was unfixed with PFA, and 52% was unfixed with Gly (**Fig 2B** and **B2**); 9% of vimentin was unfixed with PFA, and 54% was unfixed with Gly (**Fig 2C** and **C2**); 0.4% of α/β-tubulin was unfixed with PFA, and 77% was unfixed with Gly (**Fig 2D** and **D2**). The fixed protein ratios are summarized in **Table 4**.

**Fig 2.**
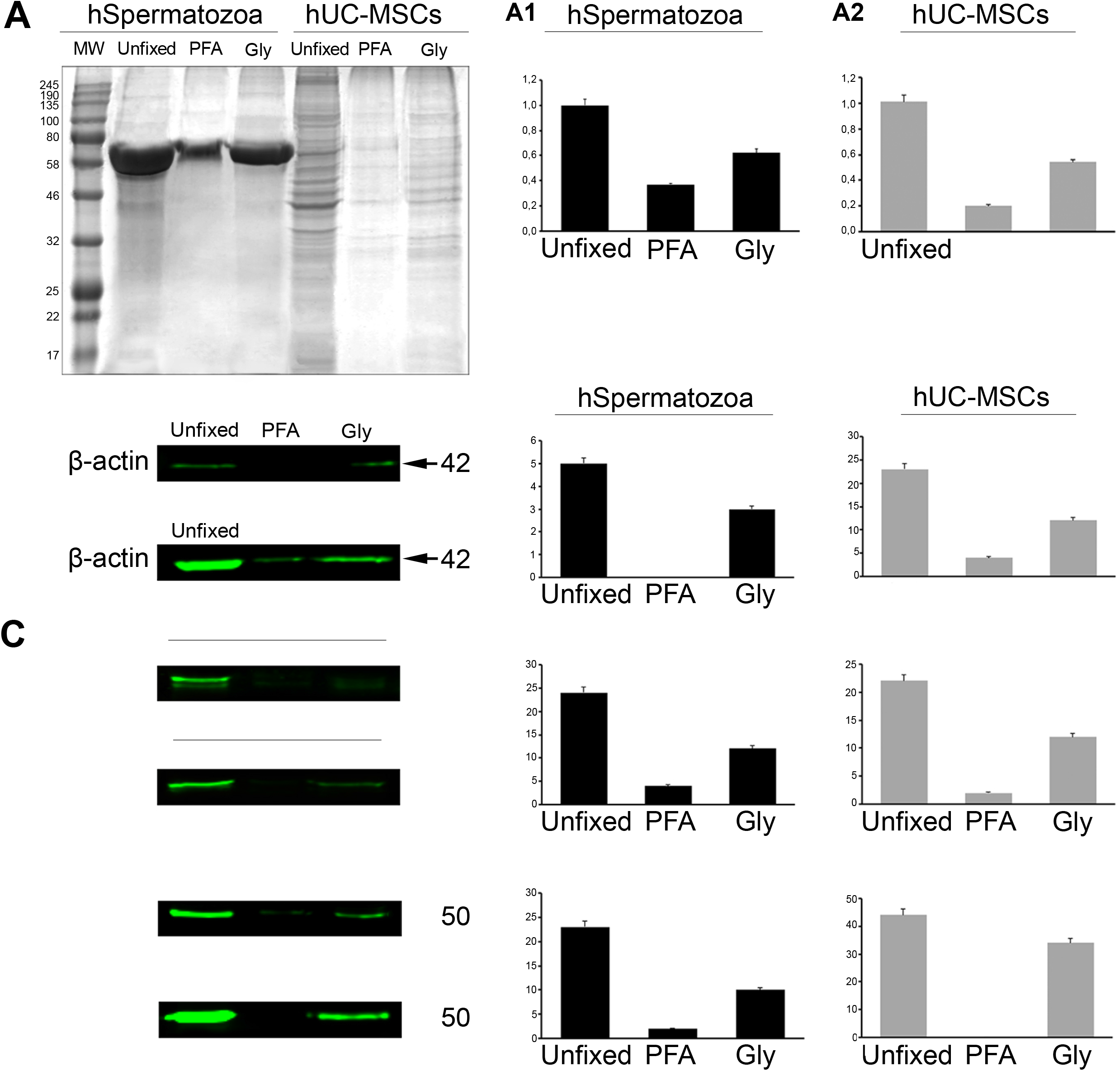
Protein fixation capacities of 3.5% PFA and 3% Gly in hSpermatozoa and hUC-MSCs. **A, A1 and A2**. Western blot assays were used to show the percentage of unfixed proteins by measuring the total signal intensity in Coomassie-blue–stained whole gel lanes in hSpermatozoa and hUC-MSC lysates; (n=6 for each group from independent experiment). **B, B1, B2**. Fluorescent-labelled 42 kD bands specific to β-actin in hSpermatozoa and hUC-MSCs. **C, C1, C2**. Fluorescent-labelled 57 kD bands specific to vimentin in hSpermatozoa and hUC-MSCs. **D, D1, D2**. Fluorescent-labelled 50 kD bands specific to α/β-tubulin in hSpermatozoa and hUC-MSCs. Data information: All intensity measurement results were found statistically different (p<0.001; Kruskall-Wallis one-way ANOVA test) from the other groups. MW: Molecular weight markers.

**Table 4.**
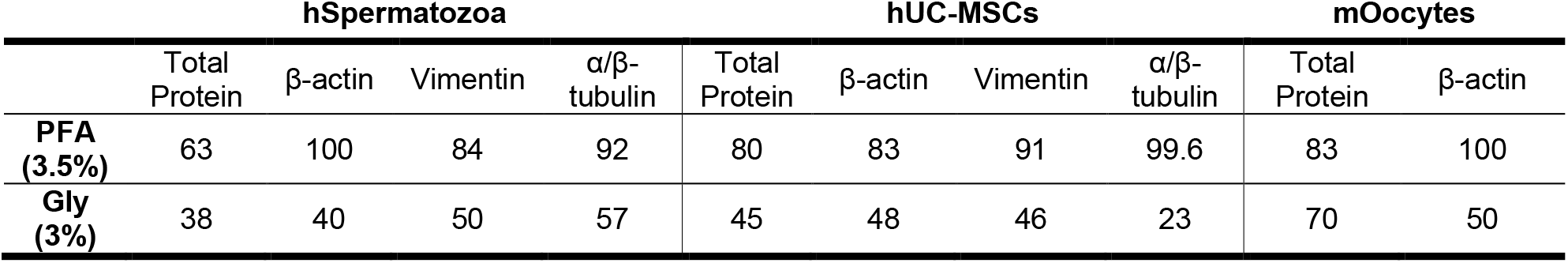
The protein fixation capacity of (% of fixed proteins) in human cell lysates after PFA and Gly fixation. The ratio of fixed vs. unfixed proteins was detected significantly higher in the PFA group (p<0.001).

As measured above, the fixation capacity of PFA and Gly was also assessed in mouse oocyte (mOocyte)/granulosa cell lysates through the evaluation of total WB Coomassie-blue stained and β-actin-labelled bands. Seventeen per cent of total protein remained unfixed in PFA, whereas 30% of proteins were found unfixed with Gly (**Fig 3A** and **A1**); 0% of β-actin was unfixed with PFA, and 50% was found unfixed with Gly (**Fig 3B** and **B1**).

**Fig 3.**
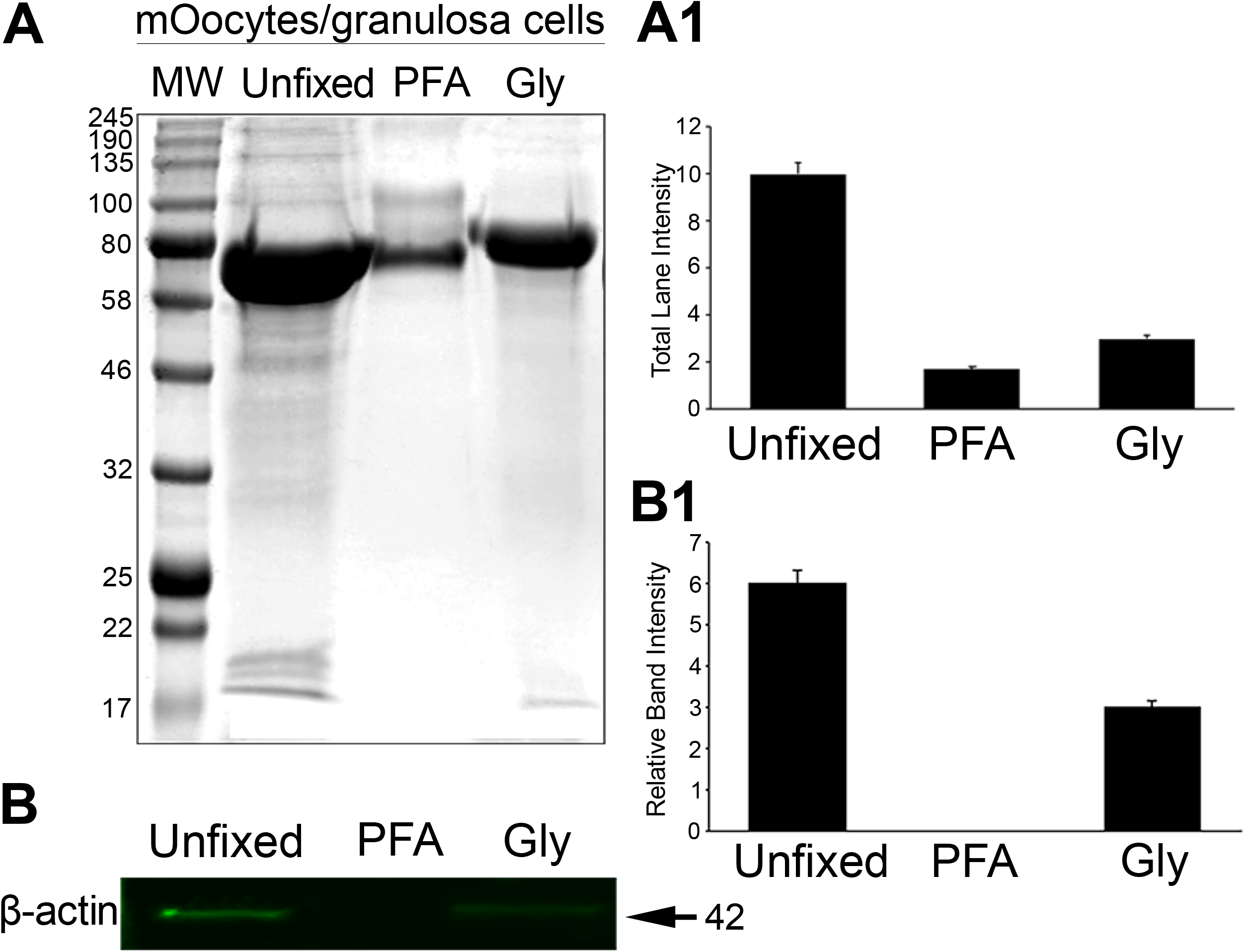
Protein fixation capacities of 3.5% PFA and 3% Gly in mOocytes/granulosa cells. **A** and **A1**. Western blot assays were used to show the percentage of unfixed proteins by measuring the total signal intensity in Coomassie-blue–stained whole gel lanes in mOocytes/granulosa cell lysates; (n=5 for each group from independent experiment). **B** and **B1**. Fluorescent-labelled 42 kD bands specific to β-actin in mOocytes/granulosa cells. Data information: All intensity measurement results were found statistically different (p<0.001; Kruskal Wallis one-way ANOVA test) from the other groups. MW: Molecular weight markers.

### 3) Qualitative and Quantitative Evaluation of Protein Labelling

#### Live-cell mitochondrial staining

Mitotracker^™^ labelling of living cells revealed fine identification of numerous mitochondria as punctate or rod-like patterns distributed throughout the entire cytoplasm of hUC-MSCs (**Fig 4**).
Following live-cell Mitotracker^™^ labelling, cells were taken into fixative cocktails after a brief wash. PFA fixation alone provided smaller numbers and lesser mitochondrial staining showing a dull appearance within the cytoplasm (**Fig 4**). In the PFA/TX group, almost no staining was observed. In contrast, significantly brighter and clearer signals were detected in the PFA+TX group. Gly fixation alone revealed high intensity and clear signals coming from mitochondria but higher background staining as well. Like in the PFA/TX groups, Gly/TX fixation provided no mitochondrial staining. In the Gly+TX group, no mitochondrial staining was observed but there was a relatively stronger background.
Live Mitotracker^™^-labelled highly motile hSpermatozoa gave positive signals restricted to their neck regions (arrows in **Fig 4**). No background or non-specific staining was observed in any of the fixatives. The brightest signals were detected in PFA alone group. The rest of the fixatives presented slightly dimmer signal intensity. TX addition to fixative did not enhance the signal intensity.

**Fig 4.**
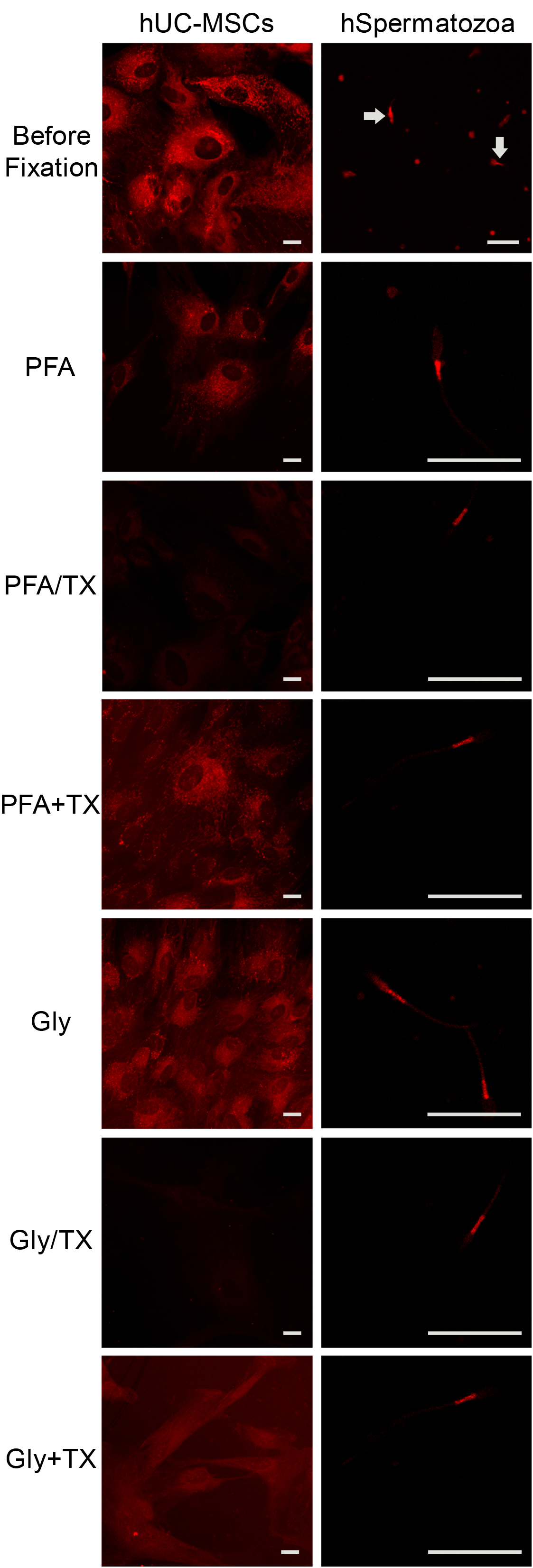
MitoTracker^™^ labelling of hUC-MSCs, hSpermatozoa after fixed with six fixative cocktails. **First row**. Live MitoTracker™ labelling of hUC-MSCs and hSpermatozoa; (n=30 labelling experiment). **Second-seventh rows**. The efficiency of fixatives on MitoTracker labeling. The best signal-to-noise ratio was noticed in PFA+TX group in hUC-MSCs. No labelling was detected in the PFA/TX and Gly/TX groups. The Gly+TX group did not display any label, but a strong background. Mitotracker™ label was confined to the neck regions of hSpermatozoa (arrows). The brightest signals were detected in the PFA group. The rest of the fixatives presented slightly dimmer signal intensity (n=5 for each group from independent labelling experiment). Data information: Scale bars: 20 μm.

#### Nuclear proteins and DNA markers

The fluorescent signal intensity and patterns of selected nuclear proteins (CENP-A, nucleostemin and lamin A/C), some of which are specific to stem/progenitor cells (i.e. nucleostemin), were tested after six different fixation cocktails in hUC-MSCs. We also examined the well-known DNA markers (Hoechst 33342 and 7-AAD) in the tiny hSpermatozoa nuclei after they were fixed with the fixatives. All results are summarized in **Fig 5**. PFA alone or with TX groups revealed fine nucleoplasmic dots, which seemed to correspond to centromeric nucleosome protein in interphase cells labelled by CENP-A antibody (**Fig 5**). PFA alone exhibited slightly higher non-specific background staining, whereas the PFA/TX and PFA+TX groups showed more specific, similar patterns and signal intensity levels. The Gly alone group also displayed punctate staining confined to the nucleus; however, a stronger cytoplasmic and nucleoplasmic background was also noted. In Gly/TX, and more commonly in the Gly+TX groups, nuclear/nucleolar nonspecific staining of irregular foci were noted (**Fig 5**, arrowheads). In the Gly+TX group, a significantly smaller number of CENP-A loci were labelled with an intense cytoplasmic background where false-positive results in signal intensity measurements were also noted.
Nucleostemin, a multilocular nucleolar protein, which was previously illustrated in hUC-MSCs (Oktar, Yildirim et al., 2011) were clearly and strongly labelled in PFA alone or PFA+TX groups in hUC-MSCs (**Fig 5**). A significantly weaker staining was noted in the PFA/TX group. Gly alone or Gly/TX groups revealed very weak signals compared with the PFA groups, whereas the Gly+TX group displayed stronger and more specific nucleostemin signals compared with the other two Gly groups.
Lamin A/C, a complex inner nuclear membrane intermediate filament protein exhibited a fairly homogenous pattern that was restricted to nuclei after hUC-MSCs were fixed with six fixatives (**Fig 5**). TX addition either to PFA or Gly showed stronger signals that were finely confined to nuclear membrane, whereas Gly alone group exhibited an extremely devastated lamin A/C pattern.
Finally, DNA dyes such as Hoechst 33342 and 7-AAD were examined in hSpermatozoa smears. Both dyes were restricted to the sperm nucleus in the head region, showing varying intensities between the fixatives. In Hoechst 33342 staining (**Fig 5**), the Gly alone group exhibited the weakest signalling intensity. 7-AAD (**Fig 5**) also displayed consistency in signal intensity; a remarkably higher signal level was noted in the Gly+TX group.

**Fig 5.**
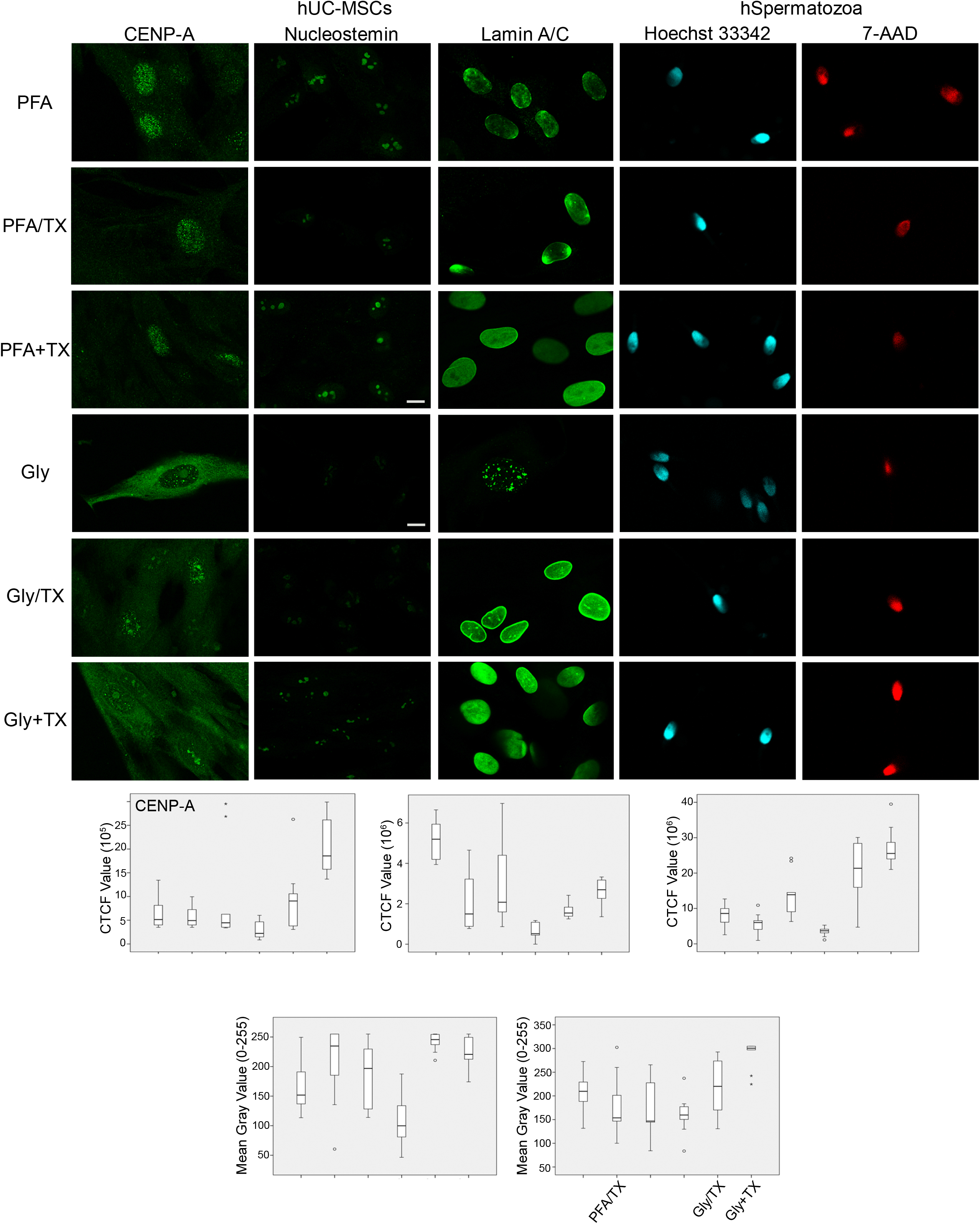
Nuclear proteins and DNA markers in hUC-MSCs and hSpermatozoa after fixed with six fixative solutions. **First column**. CENP-A were characterised by nucleoplasmic spots when fixed with PFA alone, PFA/TX, or PFA+TX. The signal intensity levels between the three groups were non-significant (p=0.999) as quantified using CTCF measurements given in the graphs. Gly alone or Gly with TX groups generally displayed intense cytoplasmic and/or nucleoplasmic background staining as well as irregular nuclear/nucleolar foci (**arrowheads**); (n=5 for labelling; n=50 for signal quantification). **Second column**. Nucleostemin in hUC-MSCs was specific but with varying intensities with no background. The highest intensity was detected in the PFA alone group; the Gly alone Gly/TX groups showed the lowest intensities (p=0.019); (n=6 for labelling; n=60 for signal quantification). **Third column**. Lamin A/C appeared predominantly on the nuclear membrane except in the Gly alone group, which displayed a disrupted arrangement. Labelling of DNA with common dyes was tested in hSpermatozoa smears as shown in the fourth and fifth columns (p<0.001); (n=5 for labelling; n=50 for signal quantification). **Fourth column**. Hoechst 33342 were consistently confined to the sperm nucleus in the head region showing varying intensities between fixatives. The Gly alone group gave the weakest signalling intensity, the difference between the other groups was insignificant (p=0.642); (n=8 for labelling; n=80 for signal quantification). **Fifth column**. 7-AAD labelling of DNA also displayed consistency in signal intensity with an exceptionally high in the Gly+TX group (p<0.0001); (n=7 for labelling; n=70 for signal quantification). Data information: Scale bars: 10 μm. All significance analysis was carried out by Kruskal Wallis one-way ANOVA test.

#### Cell-specific markers

In this section, we first demonstrate the qualitative and quantitative protein fixation results of six fixatives on various cell-specific markers tested in hUC-MSCs and hSpermatozoa. N-Cadherin, a cell adhesion molecule found primarily on the cell surface of mesenchymal cells, was selected to test the fixation capability of the fixatives, and CD73, a plasma membrane glycoprotein specific to various cell types including mesenchymal stem cells. All PFA formulations regardless of TX addition, the Gly alone and Gly+TX groups, exhibited fine, granular staining on the cell membrane without any preferential localization (**Fig 6**). N-Cadherin signals were detected as fine string-like patterns along the plasma membranes (**Fig 6**) and were clearly visualized in the PFA alone, PFA+TX, and Gly+TX groups. The remaining fixatives were unable to retain the N-Cadherin proteins. Interestingly, CD73 signals only in Gly/TX group were consistently detected both on the plasma membrane and in the cytoplasm, which may be due to the translocation of the CD73 epitopes due to the TX incubation (**Fig 6**).
Secondly, we tested the fixation potential of six fixatives on some cytoskeletal proteins (vimentin, pericentrin, and α/β-tubulin). Vimentin, an abundant intermediate filament protein, exclusively expressed in mesenchyme-originated cells, were best retained when cells were fixed with either PFA or Gly with the addition of TX (**Fig 7**). When TX was applied after fixation (PFA+TX and Gly+TX groups), the signal intensity became weak. The lowest signal was detected in the PFA alone and Gly alone groups. None of the fixatives caused any background noise or non-specific staining.
The staining of pericentrin, which is a conserved protein of the pericentriolar material, an integral component of the centrosome, was exclusively confined to two adjacent juxtanuclear foci (**Fig 7**) and best preserved in the PFA+TX and Gly+TX groups; the lowest signal intensity was measured in the Gly alone group.
As the final cytoskeletal element, microtubules composed of α- and β-tubulin were fixed with six fixatives. The results dramatically favoured PFA-containing fixatives (**Fig 7**). No significant difference was noted among the PFA-containing fixative groups. Gly-containing fixatives revealed very faint microtubule arrays (Gly alone and Gly/TX groups), and no filamentous array was noted in Gly+TX group.
Spermatozoa-specific proteins such as ACRBP, IZUMO1, and DNMT3A were examined after six fixation cocktails in hSpermatozoa (**Fig 8**). The staining pattern of ACRBP, a functional acrosin binding protein that is bound to proacrosin for the packaging and condensation of the acrosin zymogen in the acrosomal matrix, were mainly detected in the acrosomal cap region and mid-piece of spermatozoa. The strongest signal was noted in PFA/TX, Gly and Gly/TX groups, whereas the relatively weakest signal was detected in the PFA alone group.
IZUMO1, a sperm-egg fusion protein 1 that binds to its egg receptor counterpart, Juno, to facilitate recognition and fusion of the gametes, was examined after six different fixation cocktails in hSpermatozoa (**Fig 8**). The localization of IZUMO1 was found quite diverse from the acrosomal cap region to the mid-piece and neck. Signal intensity was noted as highest in the PFA group, and the weakest signal was detected in the Gly/TX group.
The signals coming from DNMT3A, an enzyme that catalyses the transfer of methyl groups to specific CpG structures in DNA, a process called DNA methylation, was strictly confined to the mid-piece (**Fig 8**). The highest signal intensity was noticed in the PFA group, and the weakest was noted in the PFA-TX group.

**Fig 6.**
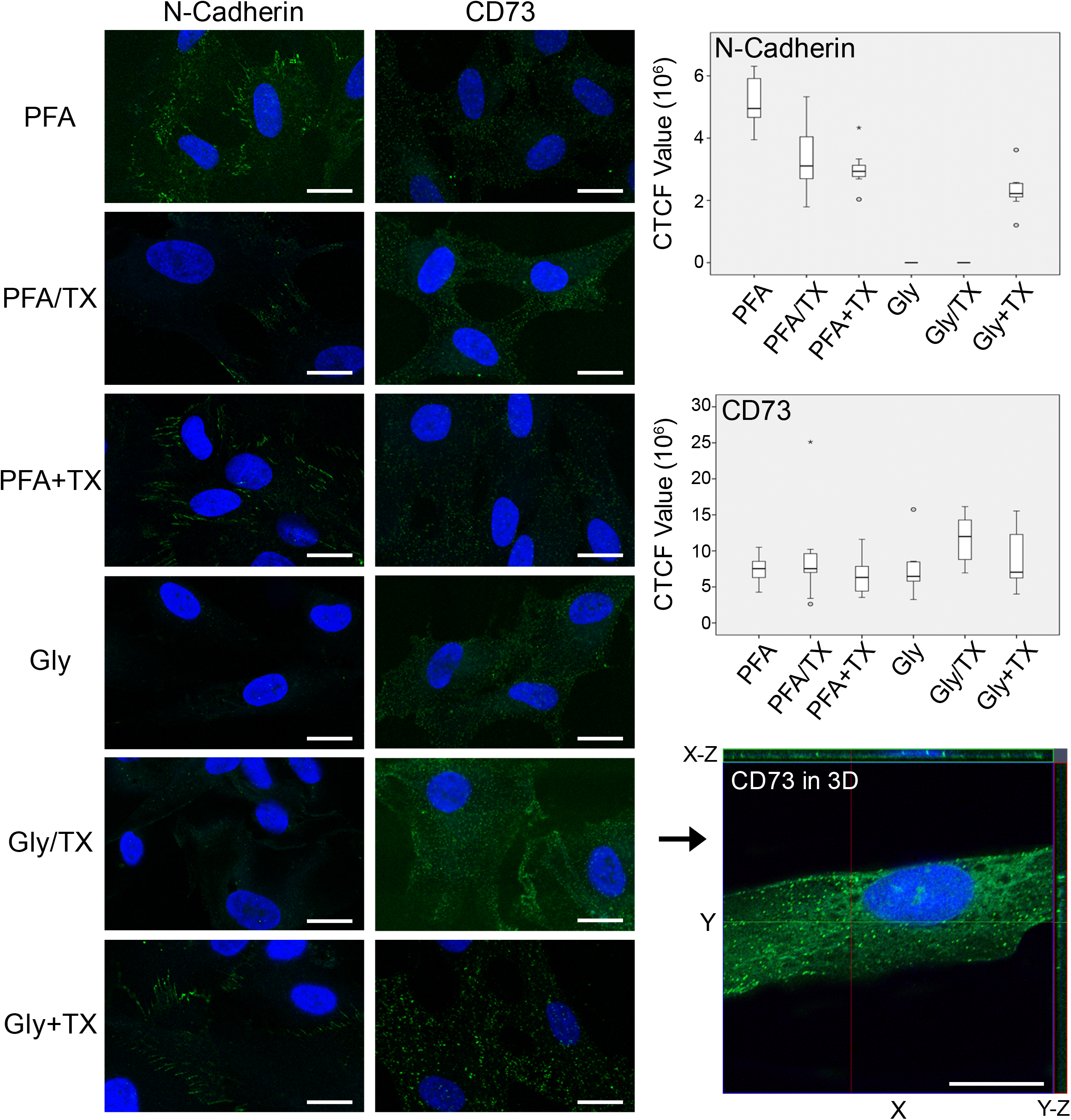
N-Cadherin and CD73 in hUC-MSCs after they were fixed with six fixative solutions. **First column**. N-Cadherin was well preserved along the plasma membrane in the PFA alone (highest compared with others; p=0.042), PFA+TX, and Gly+TX groups; (n=8 for labelling; n=80 for signal quantification). **Second column**. CD73 was well preserved in all fixative groups; however, in the Gly/TX group the signal intensity was detected significantly higher than in the other groups (p=0.021). CD73 signals in the Gly/TX group were further analysed in 3-D reconstructed image stacks obtained using super-resolution microscopy in the Z-axis (2.44 μm in thickness). X-Y axis lateral and X-Z, Y-Z axis orthogonal sections showed that most of the CD73 epitopes translocated to the cytoplasm and nucleoplasm; (n=6 for labelling; n=60 for signal quantification). Data information: Scale bars: 20 μm. All significance analysis was carried out by Kruskal Wallis one-way ANOVA test.

### 4. Surface Topology

#### hUC-MSCs

F-actin filaments, known as stress fibres in cultured cells, are commonly labelled with F-actin-specific mushroom toxins such as phallotoxins. Thus, we wanted to examine whether any signalling intensity and localization difference existed due to the different fixatives in extremely flat hUC-MSC monolayers. Fine fixation is essential for the preservation of those tiny cellular processes because cells possess wide lamellipodia and microspikes. PFA exhibited finer fixation as evidenced by the preservation of F-actin-labelled microspikes (**Fig 9**) compared with Gly, with which no microspikes were preserved.

**Fig 7.**
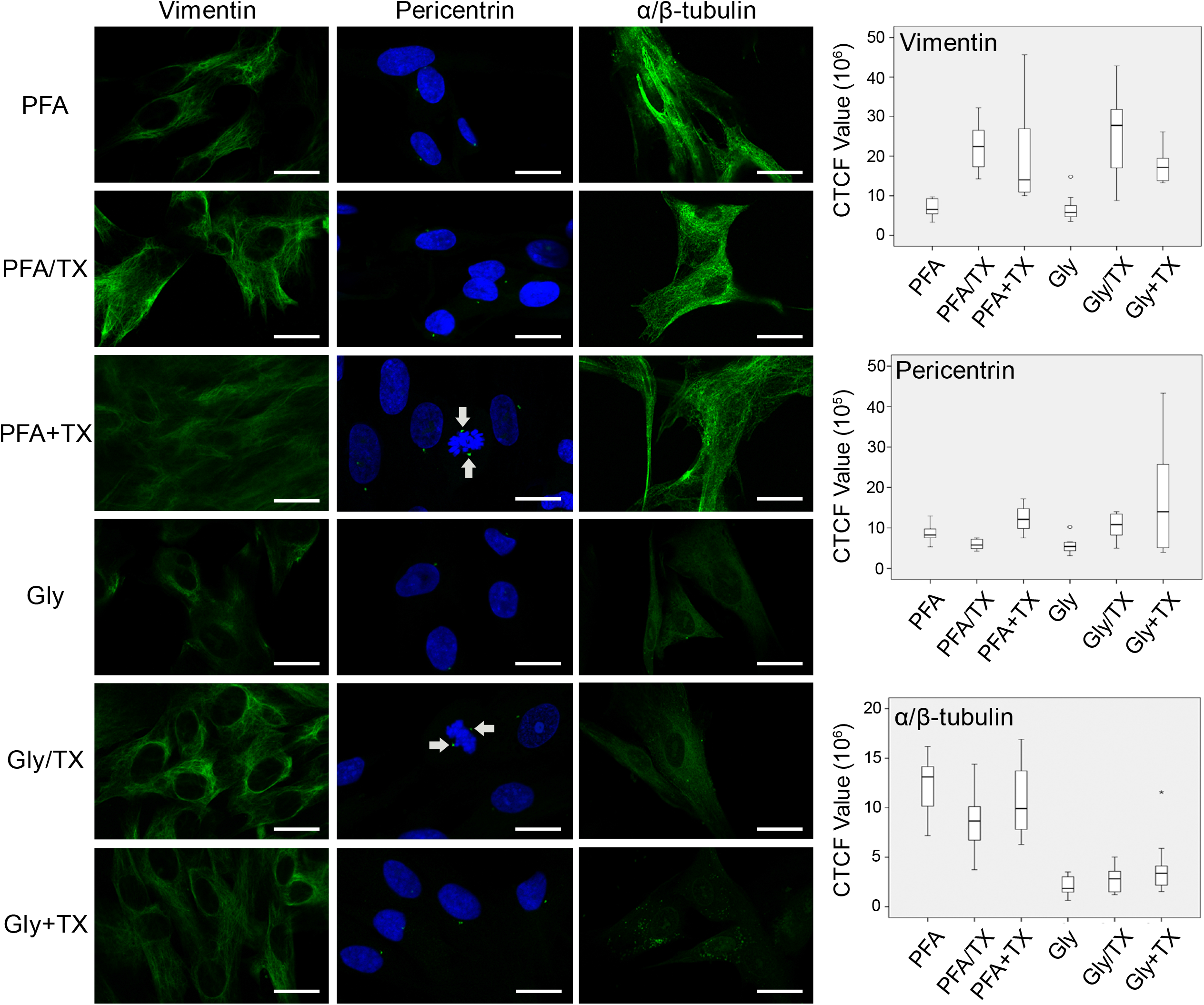
Preservation of vimentin, pericentrin, α/β-tubulin in hUC-MSCs after fixing with six fixative solutions. **First column**. Vimentin was well retained with all fixatives; however, the best and most consistent signals were found in the PFA/TX and Gly/TX groups (p=0.001); the PFA+TX and Gly groups displayed weak vimentin signals (p<0.001); (n=5 for labelling; n=50 for signal quantification). **Second column**. Pericentrin was best retained in the PFA+TX and Gly+TX groups (p=0.005); the Gly alone group revealed the lowest signal intensity (p=0.002). Pericentrin proteins were also well-preserved in occasionally encountered mitotic cells (arrows); (n=5 for labelling; n=50 for signal quantification). **Third column**. α/β-tubulin staining was best-preserved in all PFA formulations (p<0.001), whereas Gly groups in general exhibited vary faint staining. Gly+TX did not preserve any α/β-tubulin; (n=6 for labelling; n=60 for signal quantification). Data information: Scale bars: 20 μm. All significance analysis was carried out by Kruskal Wallis one-way ANOVA test.

**Fig 8.**
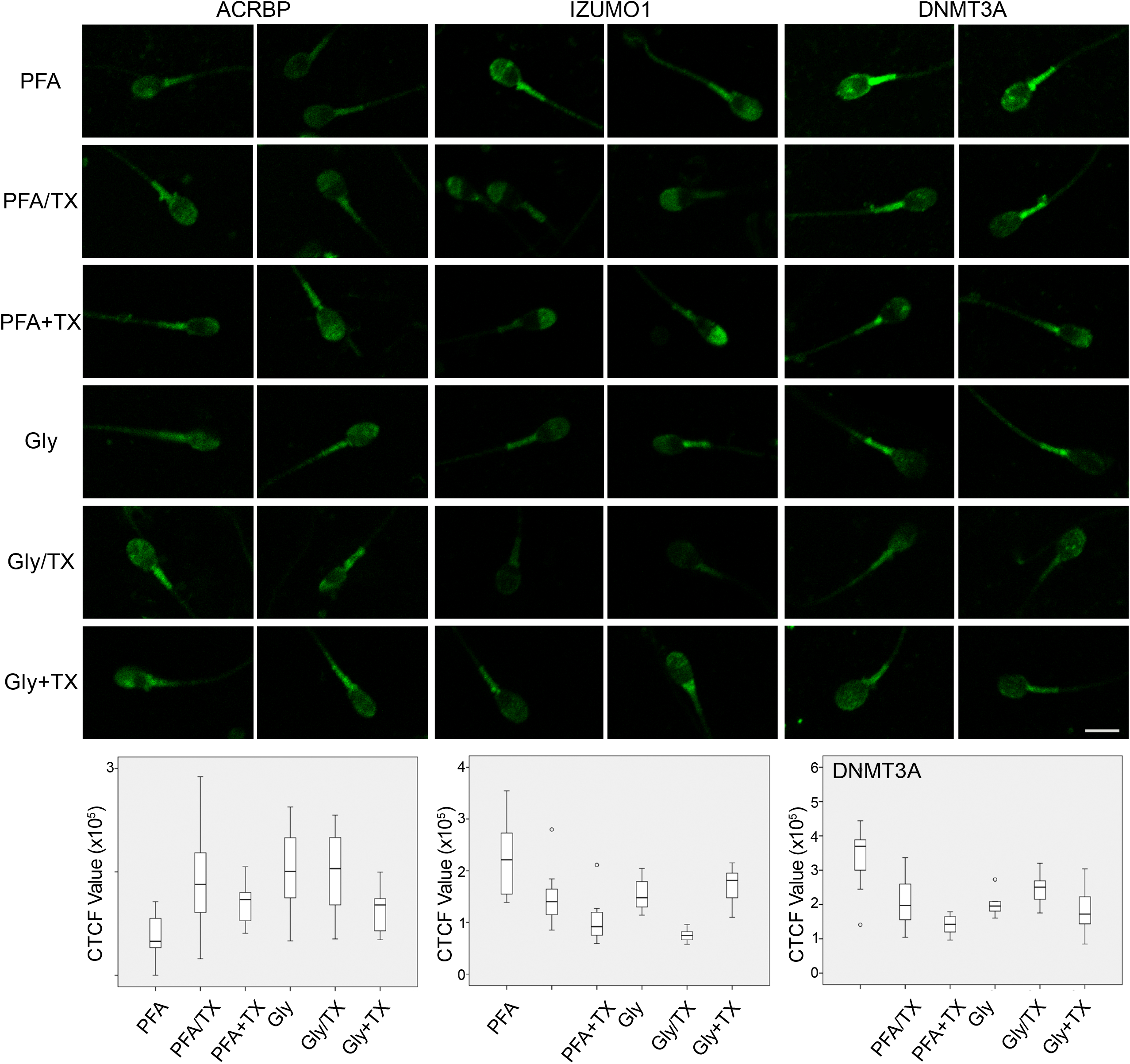
Expression of ACRBP, IZUMO1, and DNMT3A in human spermatozoa after fixing with six fixative solutions. Two set of images were provided for each fixative because all those three protein labels presented varying patterns and localizations. **First and second columns**. The ACRBP protein was mainly detected in the acrosomal cap region and mid-piece. The highest signal was noted in the PFA/TX, Gly and Gly/TX groups (p=0.004); the relatively weakest signal was detected in the PFA alone group (p=0.001) (n=6 for labelling; n=60 for signal quantification). **Third and fourth columns**. The localization of IZUMO1 showed variations from the acrosomal cap region to the mid-piece and neck. Signal intensity was noted highest in the PFA group (p<0.0001); the weakest signal was detected in the Gly/TX group (p=0.001) (n=7 for labelling; n=70 for signal quantification). **Fifth and sixth columns**. DNMT3A was rather confined to the mid piece. The highest signal intensity was noticed in the PFA group (p=0.014); the weakest was noted in the PFA-TX group (p<0.0001); (n=6 for labelling; n=60 for signal quantification). Data information: Scale bars: 5 μm. All significance analysis was carried out by Kruskal Wallis one-way ANOVA test.

**Fig 9.**
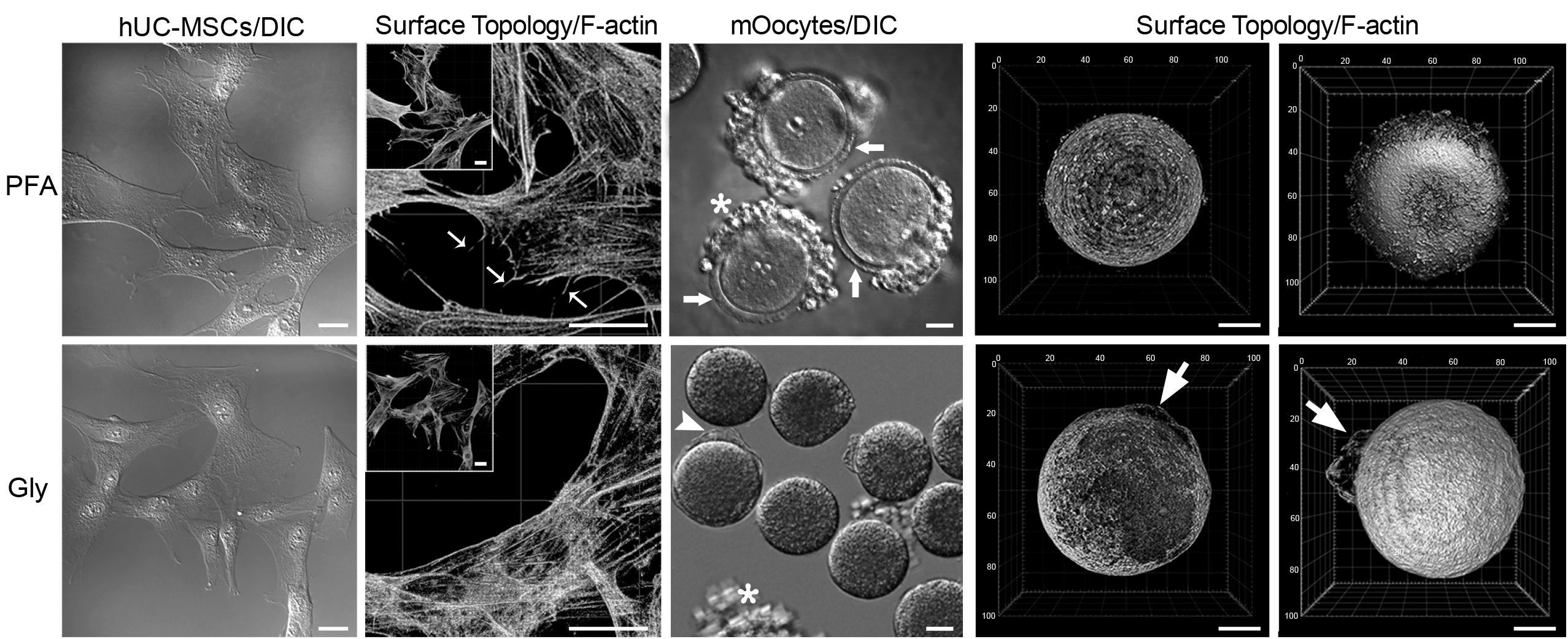
Preservation of cellular processes and plasma membranes in hUC-MSCs, and mOocytes, respectively. **First column**. DIC images showed the extremely flat, huge cell bodies of hUC-MSCs. **Second column**. F-actin-labelled microspikes (arrows) were detected by surface topological renderings. As indicated by thin arrows, microspikes were well preserved by PFA, whereas Gly was not able to retain those cellular processes; (n=3 for labelling). **Third column**. DIC images of mOocytes and cumulus-enclosed (asterisks) oocytes showed the size difference, dissolution of ZP (arrowheads) in the Gly group compared with the intact ones in PFA group (arrows); (n=16 from each independent experiment). **Fourth and fifth columns**. Surface topologic renderings confirmed the size difference, ZP dissolution, and oolemma destruction (**arrows**). Data information: Scale bars: 20 μm.

#### mOocytes

During fixation procedures with different fixatives, we were interested to note the dramatic difference between PFA and Gly fixatives regarding the morphology of naked mOocytes and cumulus enclosed oocytes. After fixing them with either PFA or Gly and then labelling with FITC-conjugated phalloidin toxins, consecutive optical sections obtained using confocal microscope allowed us to visualize those cells as three-dimensional (3D) image stacks. DIC images simply presented the size difference between two fixatives (**Fig 9**); Gly-fixed cells were significantly smaller than PFA-fixed ones. As clearly presented in **Fig 1** and the **Supplementary movie** (Gly-mEmbryos.wmv), ZP was heavily affected by Gly fixation, whereas it was intact in the PFA group. Surface rendering of image stacks showed a dramatic difference because the plasma membranes of mOocytes (oolemma) were found damaged (**Fig 9**).

## Discussion

Gly, which was first presented in 1943 by Wicks and Suntzeff, was proposed to be more efficient than 10% formalin (3.7% FA solution) and was reported to be less harmful by inhalation than FA (Sabatini et al., 1963, Wicks & Suntzeff, 1943). Thus, replacing Gly with FA in the chemical fixation of cells and tissues may be reasonable. However, studies comparing the fixation efficiency of FA with Gly in the preservation of different cellular targets (membrane receptors, cytoplasmic and nuclear targets), as well as the general morphology in tissue sections presented that FA was superior to Gly because Gly frequently resulted in significantly weaker and/or nonspecific staining (Atkins et al., 2004, Titford & Horenstein, 2005, Umlas & Tulecke, 2004).

The use of Gly as a fixative in the literature is rather old and rare compared with the other aldehyde fixatives. From the time when the fixation efficacy of PFA was recognized as better for electron microscopy in the 1970s (Smit, Meijer et al., 1974) and then for immunocytochemistry (ICC) in the 1980s, it has become the most commonly used aldehyde fixative, especially in precise microscopic observations in cell and molecular biology. In this study, we wanted to compare those two aldehyde fixatives side-by-side using super-resolution confocal microscopy to determine whether Gly, as a historic and lesser-known compound, has any superior capacity for fixation over cellular proteins. Very recently, Richter et al. published a comprehensive and well-designed data series pertaining to Gly vs. PFA fixation and concluded that 51 cellular targets were better preserved after Gly fixation (Richter et al., 2018). This led us to test Gly for our samples, most of which are related to male and female reproductive organs, early embryonic and adult stem cells. Our overall comparison of PFA and Gly results are summarized in **Table 5**, where positive and negative outcomes are designated as green and red colours, respectively.

**Table 5.**
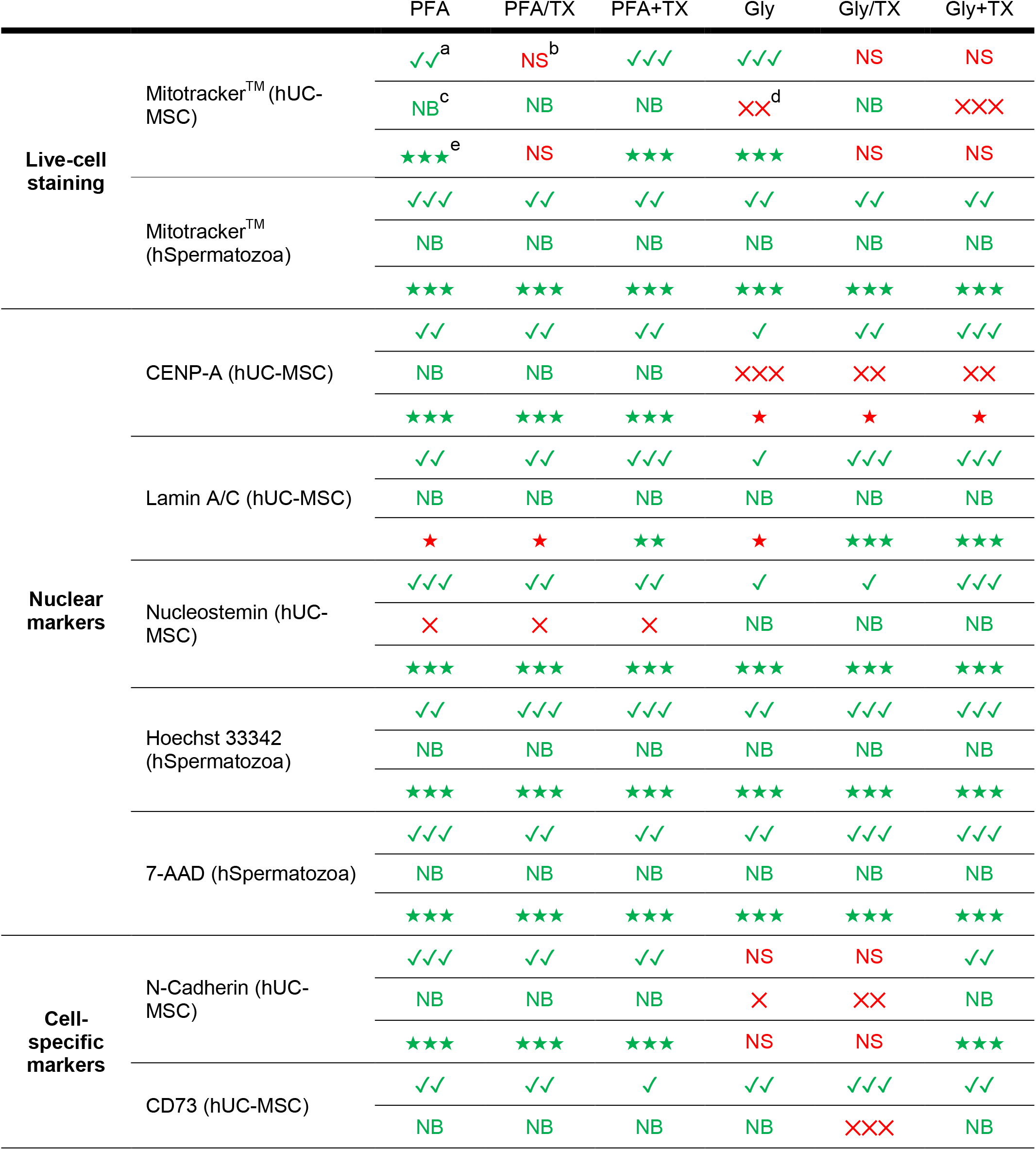
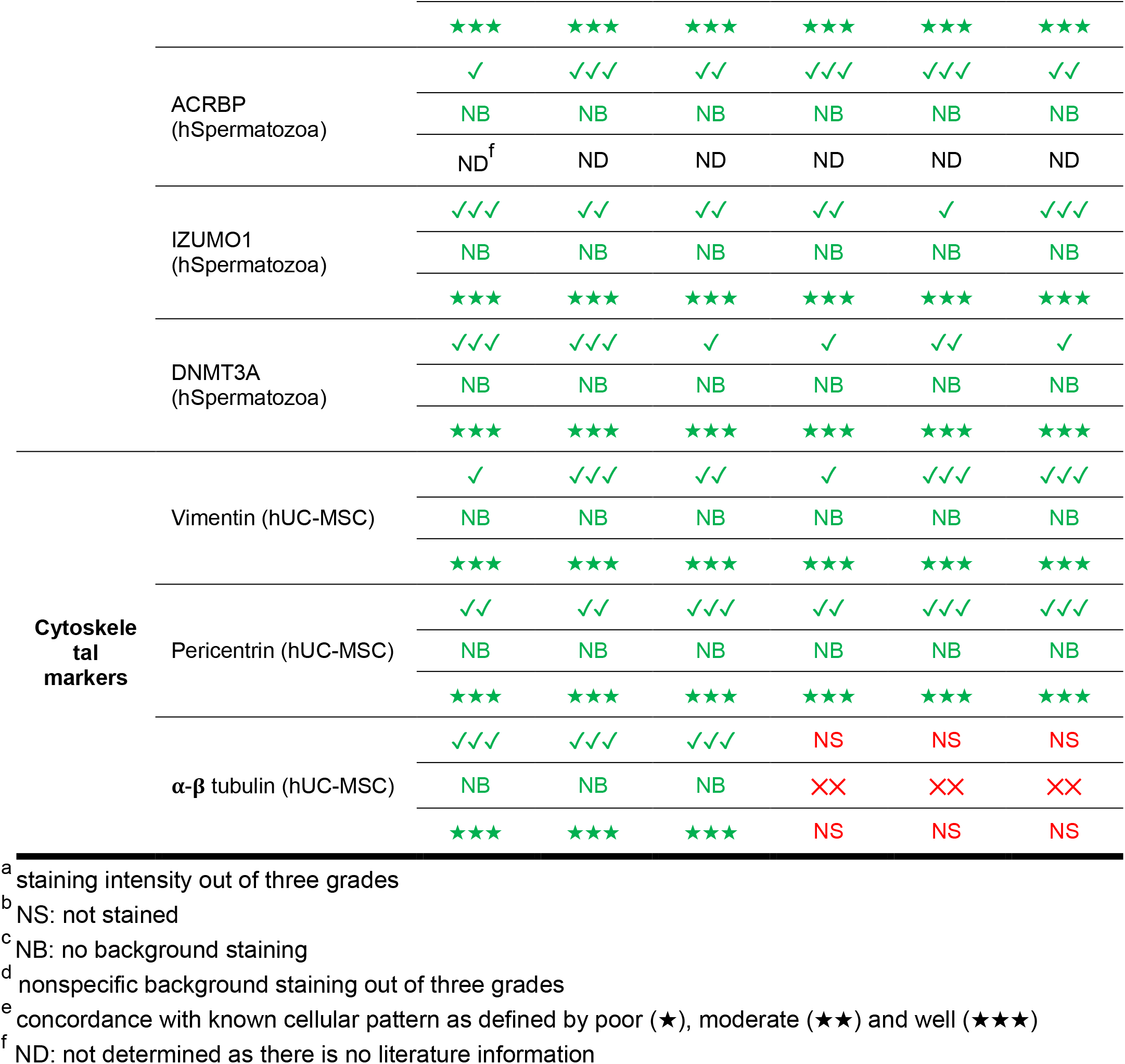
The overall performance of tested fixatives on various cell markers based on the intensity measurements and qualitative analyses using ICC. Green designates positive outcomes and red refers to negative outcomes.

In the study by Richter et al., the final pH of Gly was determined as 5 after they obtained similar results at pH 4 and 5 for most of their experiments. Therefore, we adopted Gly at pH 5 for all sets of our experiments. They calculated the proportion of the proteins that remained unfixed in brain lysates and found that 40% and 20% of proteins were unfixed with PFA and Gly, respectively. In contrast, as summarized in **Table 4**, we found that significantly more proteins remained unfixed after Gly fixation. Interestingly, Gly very consistently resulted in higher band intensities than those in PFA ones in hSpermatozoa, hUC-MSCs and mOocyte lysates where fixed proteins either no longer ran into the gel or formed smears only. More specifically, the fixation efficiency of PFA favoured Gly because β-actin, vimentin, and α/β-tubulin proteins were retained with greater proportions after PFA. In addition to the degree of β-actin preservation, we also demonstrated the lack of F-actin-decorated microspikes in Gly-fixed stem cells and surface disruptions in F-actin-labelled oocyte cortex as rendered using high-resolution consecutive fluorescent images. In contrast to our findings, different laboratories that contributed to Richter et al.’s study reported that a relatively higher ratio of β-actin, α- and β-tubulin, and F-actin was preserved with Gly. Nevertheless, they reported that a higher vimentin ratio was retained with PFA based on their ICC staining.

Live-cell imaging of embryos during fixation revealed that Gly caused thinning and then rupture of ZP, and dissociation of blastomeres, whereas tiny infrequent membrane blebs were noted in somatic cells. PFA, on the other hand, maintained the blastocyst morphology as intact as possible while generating the formation of large cytoplasmic blebs in somatic cells. Gly with TX showed similar findings in embryos while maintaining the cytoplasmic and membrane structures in somatic cells. TX application after fixation revealed no further change either in embryos or somatic cells. Based on our live-hUC-MSC videos during fixation, we agree with the statement proposed by Richter et al.,(Richter et al., 2018) that a higher speed of membrane penetration and the sudden interruption of vesicle trafficking was seen with Gly. In contrast to PFA, which resulted in the formation of large membrane blebs and continued almost during the entire fixation period, Gly displayed a very fast and relatively intact vesicle preservation with very few tiny blebs during fixation. However, Gly caused a significant decrease in the size of embryos and oocytes, as demonstrated in **Fig 1** and **Fig 9**. Although some of the above features may seem advantageous to Gly, the catastrophic changes in the embryo caused by Gly were not acceptable. We assume that Gly-originated perturbations may be due to its acidic nature, which arises from its fast oxidation, and therefore leads to the formation of strong acids, mainly glyoxylic acid (Zhang, Zhao et al., 2013).

*In situ* labelling of diverse proteins using ICC after PFA or Gly fixation did not exhibit results superior to Gly. Briefly, Gly caused a higher number of unstained epitopes, a stronger background, and nonspecific staining. Gly was occasionally found better than PFA in terms of fluorescence signal intensity.

The comparison of PFA with Gly in fixing some of the spermatozoa-specific proteins in humans such as IZUMO1 and ACRBP was first documented in this article. Consistent with its functional role, IZUMO1 was previously reported to localize in the acrosome, mid-piece, and neck regions when fixed with 3% PFA (Cakar, Cetinkaya et al., 2016). In this study, those precise localizations were confirmed, mostly with the PFA-fixed groups. The Gly-fixed groups displayed relatively weak or no staining. ACRBP in human spermatozoa has not been previously reported in the literature. In mice, ACRBP was restricted to the acrosomal region (Kanemori, Koga et al., 2016), although we consistently detected it both in the acrosomal region and the mid-piece. No significant localization or signal intensity difference was noted between the PFA and Gly groups. Likewise, DNMA3A enzyme was found localized to the mid-piece, as published previously in a single study by Marques et al. (Marques, Joao Pinho et al., 2011). Although PFA-fixed DNMT3A epitopes showed the greatest intensity, the results of all PFA and Gly-fixed samples confirmed the cellular localization published in their study.

All cross-linking reagents (all aldehyde group fixatives) form intermolecular bridges, normally through free amino groups, thus creating a network of linked antigens. Cross-linkers preserve cell structure better than organic solvents, but may reduce the antigenicity of some cell components, and require the addition of a permeabilisation step to allow access of the antibody to the specimen (Ramos-Vara & Miller, 2014). Non-ionic detergents such as TX are widely used to permeabilise and solubilise membrane proteins in a gentler manner, allowing the solubilised proteins to retain a native subunit structure and enzymatic activity (Bhairi S. M., 2001). Nonionic detergents are considered non-denaturating because they break lipid-lipid and lipid-protein, but not protein-protein interactions (Fonfría V.L., 2011); As such, they are considered for use in the isolation of membrane proteins in cell and molecular biologic applications (Bhairi S. M., 2001). Due to our TX-containing fixative results, some protein labels were partially/completely lost (i.e. Mitotracker, CD73, N-Cadherin and DNMT3A), whereas the intensity of some labels was enhanced (i.e. CENP-A, lamin A/C, nucleostemin, DNA stains, ACRBP, and vimentin).

Conclusively, the superiority of PFA over Gly was clear, at least for certain cell types and proteins as we have presented here. However, no single type of fixative cocktail and protocol seems to fit to all types of samples and proteins. Thus, it is still noteworthy to suggest that the proper fixative (with or without TX addition) formulae and fixation procedures should be carefully determined through a series of well-designed comparative experiments to optimize protein preservation as naturally as possible.

## Materials and Methods

### Preparation of PFA Stock (10%) and Working (3.5%) Solutions

Stock PFA (10%, w/v) solution was prepared by adding 100 mg PFA (95% powder) (Merck, Germany) to 650 mL double distilled water (ddH_2_O), then heated to 60°C using a magnetic heater/stirrer. Fifty microliters of 1 N NaOH was added to dissolve the PFA completely. After removing from the heater, the suspension was finalized to 1000 mL using ddH_2_O; the pH was brought to 7.2 using 1 N HCl. Stock PFA solution was filtered and cooled down to 4°C for storage. Immediately before use, to prepare 3.5% PFA working solution, stock PFA solution was mixed with ddH_2_O in 1:1.85 ratio (pH= 7.2-7.3).

### Preparation of Gly Working (3%) Solution

For preparing a 1000 mL 3% Gly working solution, 197.25 mL absolute ethanol (Merck, Germany), 79.50 mL 40% Gly (v/v) (Merck, Germany), and 7.5 mL glacial acetic acid (Merck, Germany) were added to 709.75 mL ddH_2_O. The solution was vortexed and 1N NaOH (approximately 6.20 mL 1N NaOH) were added until pH=5 has been reached. The clear solution was stored at 4°C and used within few days with no precipitation.

### Experimental Groups

The fixation potency of 3.5% PFA and 3% Gly solutions were tested in hUC-MSCs, hSpermatozoa, mOocytes and mEmbryos with or without 0.1% Triton X-100 detergent (TX). The experiment groups and fixation protocol are summarized in **Table 1**.

### Collection and Preparation of Human Cells and Mouse Oocytes/Embryos

Fresh umbilical cords (n=3) were obtained from full-term female infants after caesarean section succeeding the acquisition of informed consent from the mother (Local Ethics Committee, IRB approval number 18-578-12, 2012). The details of obtaining and culture of hUC-MSCs can be found elsewhere (Can & Balci, 2011). Briefly, umbilical cord pieces were chopped into approximately 0.5-mm^3^ tissue explants. Following the lag phase, sprouting hUC-MSCs cells were expanded in 75-cm^2^ tissue culture flasks (Corning; 430641, USA) in DMEM-Ham’s F12 media (Biochrom T481-01, Germany) at 37°C 5% CO_2_ until passage 4 (P_4_) cells reached 90-95% confluence. At the final stage, confluent cells were removed from flasks using Trypsin-EDTA (0.05/0.02%) (Biochrom, Germany) and then transferred onto poly-L-lysine–coated (0.01%) (Sigma-Aldrich, USA) glass coverslips (5000 cells/cm^2^) and left for at least 48 hr before fixation. Prior to fixation, the procedure cells were washed twice with Dulbecco’s PBS (D-PBS).

Ejaculates were obtained from undisclosed healthy donors (n=4) by masturbation after 3 days of abstinence. The experiment protocol was approved by the IRB (Local Ethics Committee approval number: 05-298-18, 12.05.2018). Informed consent was obtained from all subjects and that the experiments conformed to the principles set out in the WMA Decleration of Helsinki.

The details of preparation of hSpermazoa can be found elsewhere (Sati & Huszar, 2013). Briefly, following liquefaction, semen samples were washed in sperm wash medium (SpermRinse™, Vitrolife, Sweden) and the spermatozoa were pelleted by centrifugation at 800 *g* for 20 min at room temperature (RT). Following centrifugation, the pellet was resuspended in sperm wash medium and adjusted to a final concentration of 1×10^7^ sperm/mL. The sperm slides were prepared as follows: phosphate-buffered sucrose (PB-Sucrose) pools [22.5 g sucrose (Merck, Germany) dissolved in 500 mL ddH_2_O containing 20 mL 0.5 M sodium phosphate buffer (Merck, Germany), pH: 7] were made by adding 2 or 3 drops onto each 5% poly-L-lysine–coated glass slides; then 2–5 μL of the sperm suspension were pipetted into the PB-Sucrose pool. The slides were incubated overnight (o/n) in a humidified chamber at 4°C, allowing the spermatozoa to settle and attach to the surface of the poly-L-lysine–coated slides. The following day, the spermatozoa were fixed with the fixation solutions for 15 min.

Oocytes and embryos were obtained from BALB/c female mice (n=8) at 4-6 weeks. The experimental protocol was approved by the IRB (approval number 792/20-18.10.27). The experimental protocol was approved by the Animal Care and Usage Committee (Protocol no: B.30.2.AKD.0.05.07.00/30). All mice were hosted with free access to food and water and kept in a 12 hr-light/dark cycle. The germinal vesicle (GV)-, meiosis-I (MI)-stage oocytes and zygote/blastocyst-stage embryos (n=275) were obtained from mice primed with 5 IU (0.1 mL/animal) pregnant mare’s serum gonadotropin (PMSG) (Intervet, UK). The details of the collection of mOocytes can be found elsewhere (Can, Semiz et al., 2003). For embryo collection, mice were further injected with 5 IU (0.1 mL/animal) human chorionic gonadotropin (hCG) (Sigma-Aldrich, USA). Twenty hours after PMSG injection, the mice were killed and then the ovaries were punctured with a 23-gauge needle in G-MOPS medium (Vitrolife, Sweden). GV-stage oocytes were collected using a mouth-controlled pipette under a dissecting microscope (Nikon, Japan). hCG-induced female mice were mated overnight with mature male mice at a rate of 1 female:1 male. Upon confirming a vaginal plug, zygote-blastocyst stage embryos were collected in the following days from the oviducts/uterus.

### Live-cell Imaging

hUC-MSCs (1×10^5^ cells/cm^2^) were plated onto glass-bottomed 35-mm Petri dishes (World Precision Instruments, USA) in complete medium (DMEM/HAM’s F12 + 10% FBS) (Merck, Germany), whereas live mEmbryos were cumulated in a 50 μL drops containing GMOPS covered with Ovoil^™^ (Vitrolife, Sweden) in glass-bottomed 35-mm Petri dishes. Live hUC-MSCs and mEmbryos were viewed using live-cell imaging microscopy (controlled temp, CO_2_ and humidity) before and during the fixation period. Scoping time (min), temperature (°C) and image acquisition interval (sec) were aligned according to the cell type and fixation groups (**Table 2**).

All movies were recorded in a time-lapse manner for 15/20 min with 5 or 8-second intervals using differential interference microscopy (DIC) configured on a Zeiss LSM-880 confocal system (Zeiss, Germany) equipped with a Zeiss Axio Observer inverted microscope. A 633 nm red laser line (5 mW) was used as a light source. At the objective plane (Zeiss 20x plan apo/NA 0.8), the laser power was measured 1.97 mW. The laser power was set to 1.2% during image acquisition to keep the excitation energy as low as possible to avoid any photo damage/toxicity; therefore, the cells and embryos were scanned with 0.023 mW (23 μW). Scanning parameters were set in order to achieve pixel dwell time between 0.40-0.50 microsecond.

### Western Blotting

Total protein, β-actin, vimentin, and α/β-tubulin quantities in unfixed and fixed hUC-MSCs, and hSpermatozoa and mOocytes/granulosa cells were determined using Western blotting. hUC-MSCs and hSpermatozoa were fixed for 15 min, and mOocytes were fixed for 20 min at RT. Fixed and non-fixed cells were dissolved in lysis buffer [1% sodium dodecyl sulphate (SDS), 1.0 mM sodium ortho-vanadate, 10 mM Tris pH 7.4] supplemented with 1x protease inhibitor cocktail (Amresco, USA). The protein concentration was measured using the BCA method. Fifty micrograms of protein from each group was separated on 10% Tris-HCl gel using protein electrophoresis (BioRad, USA). The gels were stained in Coomassie brilliant blue overnight (o/n). In the following day, they were de-stained in a 50% methanol, 40% ddH_2_O, and 10% acetic acid mixture for 3-4 hr. The stained gels were scanned and analyzed as described below.

To determine the levels of the aforementioned proteins, 50 μg of cell extract from each group was separated on 10% Tris-HCl gel and then electrotransferred to a polyvinylidene difluoride (PVDF) membrane (Roche, UK) o/n at +4°C. Then, the membrane was blocked with 5% (w/v) bovine serum albumin (BSA) prepared in TBS-T (20 mM Tris/HCl and 150 mM NaCl plus 0.05% Tween-20 at pH 7.4) at RT for 1 hr. Membranes were incubated with primary antibodies specific to β-actin (Abcam; ab8226) vimentin (Sigma; V6630) or α/β-tubulin (Sigma; T9026, T4026) [1:1000 in 5% (w/v) BSA containing TBS-T] for 2 hr at RT. Following a triple- wash in TBS-T for 15 min each, membranes were incubated with goat anti-mouse or anti-rabbit secondary antibody (800 nm) (1:2000 in TBS-T) (Licor Biosciences, USA) at RT for 1 hr on a shaker. The SDS–PAGE gels in **Fig 2** and **Fig 3** were analysed by measuring the overall lane intensity that was left after fixation and compared with the non- fixed sample. Protein band intensities were measured using a Li-Cor Odyssey CLx infrared detection system (LICOR Biosciences, USA) following the manufacturer’s guidelines. Relative band intensities were measured and analysed using ImageJ v.3.91 software.

### Live-Cell Mitochondrial Staining

Live hSpermatozoa in sperm wash medium were incubated with 1 mM MitoTracker™ (Molecular Probes; M7512, USA) for 45 min at 37°C. hSpermatozoa were then washed twice with PBS and fixed with six different fixative cocktails for 15 min at RT. In PFA+TX and Gly+TX groups, 0.1% TX was applied for 5 min at RT following fixation and washing. After the final incubation step, the hSpermatozoa suspension was subjected to centrifugation for 15 min at 800 g, the cell pellet was washed with PBS, and then 50 μL mounting medium (MM) (1:1 glycerol/PBS and 0.002% NaN_3_) was added onto the cell pellet. Finally, a drop of labelled hSpermatozoa was gently laid down onto a glass slide and then covered with a glass coverslip.

### Nuclear Markers

Fixed hUC-MSCs were incubated with antibodies against nuclear proteins such as CENP-A (1:100 in PBS; o/n at 4°C + 2 hr at 37°C) (Abcam; ab45694, USA), lamin A/C (1:100 in PBS; following blocking with NGS, 2 hr at 37°C) (Abcam; ab108595, USA), or nucleostemin (1:250 in PBS; 2 hr at 37°C) (Chemicon International; MAB4311, USA). FITC-conjugated anti-rabbit IgG (Abcam; ab6717, USA) was used as a secondary antibody. For nuclear DNA labelling in hSpermatozoa, 7-AAD was used (1:100 in PBS) (Sigma; 9400) for 15 min at RT followed by mounting with MM or Hoechst 33342 (1:1000 in glycerol/PBS) (Invitrogen; LSH3570) in MM.

### Cell-specific Markers

Fixed hUC-MSCs with six different fixative cocktails were immunostained with cell-specific markers such as N-cadherin or CD73 as follows; N-Cadherin [1:100 in PBS; o/n at 4 °C + 2 hr at 37°C] (Sigma; C3865, USA), CD73 (1:100 in PBS; 2 hr at 37°C) (Abcam; ab54217, USA). Fluorescein isothiocyanate (FITC) conjugated anti-mouse IgG (Jackson ImmunoResearch; 115-095-166, USA) was used as a secondary antibody.

Fixed hSpermatozoa slides prewashed in PBS were initially blocked with normal goat serum (NGS) for 1 hr at RT. Then slides were incubated with antibodies specific to acrosin binding protein (ACRPB) (1:100 in PBS) (Sigma; HPA039081, USA), IZUMO1 (1:100 in PBS) (Sigma; HPA038104, USA) or DNMT3A (DNA cytosine-5-methyltransferase 3A protein) (1:200 in PBS) (Abcam; ab23565, USA) at 4°C o/n. Following incubation, the sections were washed three times in PBS for 5 min each, and incubated with FITC-conjugated goat anti-rabbit secondary antibody (1:100 in PBS) (Abcam; ab6717, USA) for 1 hr at 37°C. Slides were then washed in PBS for 5 min, and covered with MM.

### Cytoskeletal Markers

Fixed hUC-MSCs were incubated with antibodies against some cytoskeletal proteins such as vimentin (1:50 in PBS; o/n at 4°C + 2 hr at 37°C) (Sigma; V6630, USA), pericentrin (1:1000 in PBS; 2 hr at 37°C) (Abcam; ab4448, USA) or α/β-tubulin (1:1 mixture) (1:100 in PBS; 2 hr at 37°C) (Sigma; T9026, T4026, USA). FITC-conjugated anti-mouse IgG (Jackson ImmunoResearch; 115-095-166, USA) was used as a secondary antibody for anti-vimentin and anti-α/β-tubulin antibodies and FITC-conjugated anti-rabbit IgG (Abcam; ab6717, USA) for anti-pericentrin. The slides were covered with MM. Fixed hUC-MSCs and mOocytes were incubated with fluorescein phalloidin (ThermoFisher Scientific; F432, USA) (35 mg/mL in PBS for 60 min) for F-actin staining. All staining steps for mOocytes were performed using micro-well trays (Thermo Scientific, USA). Then, they were gently transferred onto glass-bottomed 35-mm Petri dishes in a 100 μL drop of PBS-based mounting medium containing 1μg/mL Hoechst 33342 (Invitrogen; LSH3570) for DNA labelling. The top was covered with paraffin oil. Mouse oocytes have a diameter of around 100 μm (including zona pellucida; ZP); therefore, this type of preparation enabled us to visualise the oocyte surface and the inner structures as naturally as possible.

### Fixed-cell Imaging with Super-Resolution Confocal Microscopy

All fluorescently-tagged specimens were examined and imaged using a Zeiss LSM-880 Airy Scan system (Zeiss, Germany). Multiple laser lines (405, 488, 543 and 633 nm) were used according to the fluorescent probes. All laser and scanning parameters were set identically for each protein using a reference histogram; 20x (dry) 40x (water), and 63x (oil) immersion objectives, and gallium-arsenide phosphide or airyscan detectors were used to detect signals. Pixel resolution for 2-D and 3-D images were set automatically and aligned identically for all image acquisitions. For 3-D surface topology assessment of GV-MI-stage oocytes, 3D module of Zen Desk software (v2.3) was used to analyse the app. 100 μm-thick image stacks.

### Quantification and Analysis of Fluorescent Signals

Ten sample images were obtained from each label (protein). The varying signal intensities were analysed using Zen Blue v2.3 histogram tool to calculate the corrected total cell fluorescence [CTCF= Integrated density – (area of selected cell) × (mean fluorescence of background readings)] (Coskun & Can, 2015). The total image area was used to calculate the CTCF value for N-Cadherin, CD73, CENP-A, lamin A/C, nucleostemin, pericentrin, vimentin, and α/β-tubulin. For sperm-specific markers and MitoTracker™, CTCF calculations were implemented on 100×100 pixel areas, which solely covered the sperm head and neck regions. For nuclear markers (Hoechst 33342 and 7-AAD), mean grey scale values (0-255) from a 1.5-μm-diameter circle area in the nuclear region was used to compare signal intensities between the different fixation groups.

### Statistical Analysis

All statistical analyses were performed using the IBM SPSS for Windows v20.0 software package (SPSS, Chicago, IL, USA). The Kolmogorov-Smirnov test was used to assess the assumption of normality. For non-normally distributed continuous variables, differences between groups were tested using the Kruskal-Wallis test. Continuous variables that did not have normal distribution are expressed as median (minimum-maximum). A two-sided p value<0.05 was considered as statistically significant. Data are reported as mean±standard deviation (SD). Sum, average and SD calculations were performed using MS Excel (Microsoft Corp, Seattle, USA). Significance tests were performed using SPSS 10.0 (SPSS Inc., Chicago, IL, USA). Data were analysed using one-way analysis of variance (ANOVA) and Student’s t-test when values were normally distributed; otherwise, the Mann–Whitney U test was applied. Differences between the experimental and control groups were regarded as statistically significant when p<0.05.

## Acknowledgements

This study was financially supported by The Scientific and Technological Research Council, and University Research Fund (17A0230001).

## Author contributions

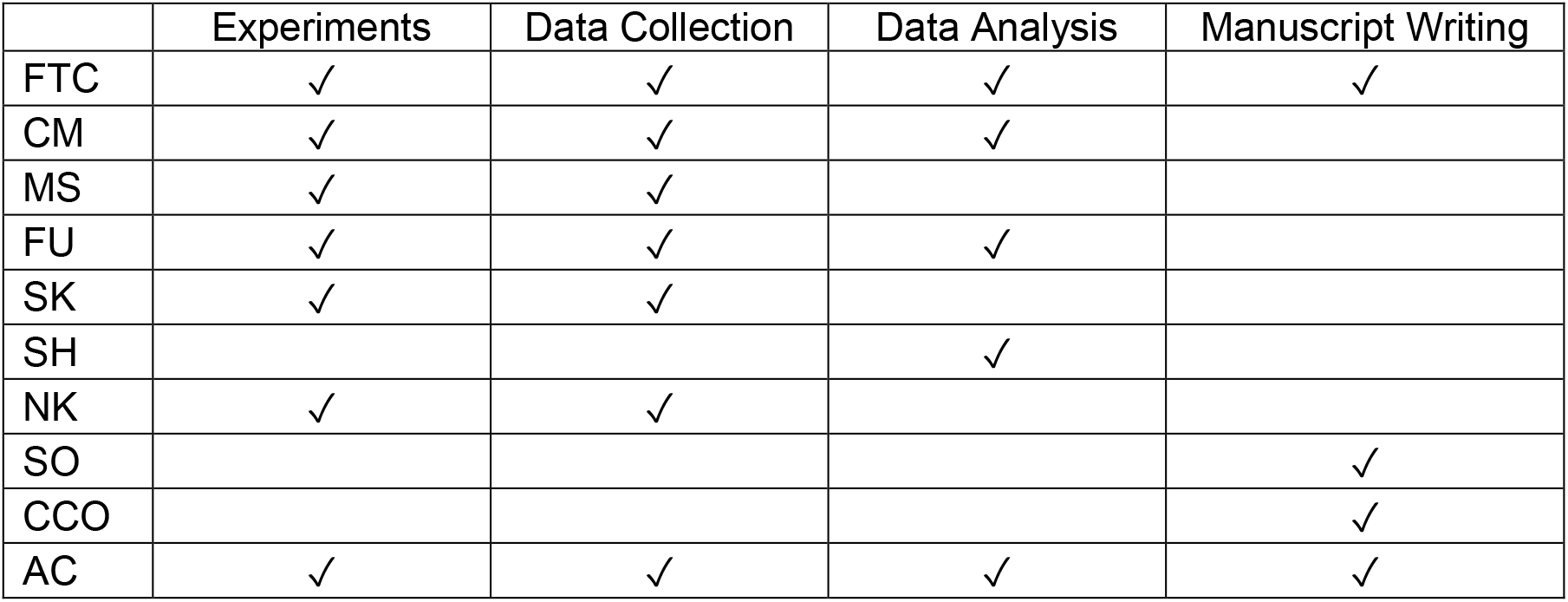

## Supplementary Movie Legends

**mEmbryos (1-7)**

Seven movie files are presented showing 20-min long time-lapse movies (40x) before and during the fixation with PFA and PFA/TX groups. Additional 5 min-TX incubation is shown in the PFA+TX and Gly+TX groups; (n=12 independent experiment).

Data information: Scale bars: 50 μm.

**hUC-MSCs (8-14)**

Seven movie files are presented showing 15-min long time-lapse movies (40x) before and during the fixation with PFA and PFA/TX groups. Additional 5 min-TX incubation is shown in PFA+TX and Gly+TX groups; (n=13 independent experiment).

Data information: Scale bars: 50 μm.

## References

Atkins D, Reiffen KA, Tegtmeier CL, Winther H, Bonato MS, Storkel S (2004) Immunohistochemical detection of EGFR in paraffin-embedded tumor tissues: variation in staining intensity due to choice of fixative and storage time of tissue sections. J Histochem Cytochem 52: 893–901

Bhairi S. M. MC, Ibryamova S., LaFavor T. (2001) Detergents: A guide to the properties and uses of detergents in biological systems. Calbiochem-Novabiochem Corporation, San Diego, CA

Bussolati G, Annaratone L, Berrino E, Miglio U, Panero M, Cupo M, Gugliotta P, Venesio T, Sapino A, Marchio C (2017) Acid-free glyoxal as a substitute of formalin for structural and molecular preservation in tissue samples. PLoS One 12: e0182965

Cakar Z, Cetinkaya B, Aras D, Koca B, Ozkavukcu S, Kaplanoglu I, Can A, Cinar O (2016) Does combining magnetic-activated cell sorting with density gradient or swim-up improve sperm selection? J Assist Reprod Genet 33: 1059–65

Can A, Balci D (2011) Isolation, culture, and characterization of human umbilical cord stroma-derived mesenchymal stem cells. Methods Mol Biol 698: 51–62

Can A, Semiz O, Cinar O (2003) Centrosome and microtubule dynamics during early stages of meiosis in mouse oocytes. Mol Hum Reprod 9: 749–56

Coskun H, Can A (2015) The assessment of the in vivo to in vitro cellular transition of human umbilical cord multipotent stromal cells. Placenta 36: 232–9

Dapson R. W. FAT, Wolfe D. (2006) Glyoxal Fixation and Its Relationship to Immunohistochemistry. Journal of Histotechnology 29: 66–76

Dapson RW (2007) Glyoxal fixation: how it works and why it only occasionally needs antigen retrieval. Biotech Histochem 82: 161–6

Dimitriadis GJ (1979) Effect of detergents on antibody-antigen interaction. Anal Biochem 98: 445–51

Fonfría V.L. Map, Padrós E., Lazarova T. (2011) Solubilization, Purification, and Characterization of Integral Membrane Proteins.

Fujiwara K (1980) Techniques for localizing contractile proteins with fluorescent antibodies. Curr Top Dev Biol 14: 271–96

Howat WJ, Wilson BA (2014) Tissue fixation and the effect of molecular fixatives on downstream staining procedures. Methods 70: 12–9

Kanemori Y, Koga Y, Sudo M, Kang W, Kashiwabara S, Ikawa M, Hasuwa H, Nagashima K, Ishikawa Y, Ogonuki N, Ogura A, Baba T (2016) Biogenesis of sperm acrosome is regulated by pre-mRNA alternative splicing of Acrbp in the mouse. Proc Natl Acad Sci U S A 113: E3696–705

Kim SO, Kim J, Okajima T, Cho NJ (2017) Mechanical properties of paraformaldehyde-treated individual cells investigated by atomic force microscopy and scanning ion conductance microscopy. Nano Converg 4: 5

Leyton-Puig D, Kedziora KM, Isogai T, van den Broek B, Jalink K, Innocenti M (2016) PFA fixation enables artifact-free super-resolution imaging of the actin cytoskeleton and associated proteins. Biol Open 5: 1001–9

Marcon N, Bressenot A, Montagne K, Bastien C, Champigneulle J, Monhoven N, Albuisson E, Plenat F (2009) [Glyoxal: a possible polyvalent substitute for formaldehyde in pathology?]. Ann Pathol 29: 460–7

Marques CJ, Joao Pinho M, Carvalho F, Bieche I, Barros A, Sousa M (2011) DNA methylation imprinting marks and DNA methyltransferase expression in human spermatogenic cell stages. Epigenetics 6: 1354–61

Oktar PA, Yildirim S, Balci D, Can A (2011) Continual expression throughout the cell cycle and downregulation upon adipogenic differentiation makes nucleostemin a vital human MSC proliferation marker. Stem Cell Rev 7: 413–24

Paavilainen L, Edvinsson A, Asplund A, Hober S, Kampf C, Ponten F, Wester K (2010) The impact of tissue fixatives on morphology and antibody-based protein profiling in tissues and cells. J Histochem Cytochem 58: 237–46

Ramos-Vara JA, Miller MA (2014) When tissue antigens and antibodies get along: revisiting the technical aspects of immunohistochemistry--the red, brown, and blue technique. Vet Pathol 51: 42–87

Richter KN, Revelo NH, Seitz KJ, Helm MS, Sarkar D, Saleeb RS, D’Este E, Eberle J, Wagner E, Vogl C, Lazaro DF, Richter F, Coy-Vergara J, Coceano G, Boyden ES, Duncan RR, Hell SW, Lauterbach MA, Lehnart SE, Moser T et al. (2018) Glyoxal as an alternative fixative to formaldehyde in immunostaining and super-resolution microscopy. EMBO J 37: 139–159

Robinson RW, Snyder JA (2004) An innovative fixative for cytoskeletal components allows high resolution in colocalization studies using immunofluorescence techniques. Histochem Cell Biol 122: 1–5

Sabatini DD, Bensch K, Barrnett RJ (1963) Cytochemistry and electron microscopy. The preservation of cellular ultrastructure and enzymatic activity by aldehyde fixation. J Cell Biol 17: 19–58

Sati L, Huszar G (2013) Methodology of aniline blue staining of chromatin and the assessment of the associated nuclear and cytoplasmic attributes in human sperm. Methods Mol Biol 927: 425–36

Smit JW, Meijer CJ, Decary F, Feltkamp-Vroom TM (1974) Paraformaldehyde fixation in immunofluorescence and immunoelectron microscopy. Preservation of tissue and cell surface membrane antigens. J Immunol Methods 6: 93–8

Titford ME, Horenstein MG (2005) Histomorphologic assessment of formalin substitute fixatives for diagnostic surgical pathology. Arch Pathol Lab Med 129: 502–6

Umlas J, Tulecke M (2004) The effects of glyoxal fixation on the histological evaluation of breast specimens. Hum Pathol 35: 1058–62

Wang YN, Lee K, Pai S, Ledoux WR (2011) Histomorphometric comparison after fixation with formaldehyde or glyoxal. Biotech Histochem 86: 359–65

Wicks LF, Suntzeff V (1943) Glyoxal, a Non-Irritating Aldehyde Suggested as Substitute for Formalin in Histological Fixations. Science 98: 204

Zhang Z, Zhao D, Xu B (2013) Analysis of glyoxal and related substances by means of high-performance liquid chromatography with refractive index detection. J Chromatogr Sci 51: 893–8

